# Characterizing macroinvertebrate community composition and abundance in freshwater tidal wetlands of the Sacramento-San Joaquin Delta

**DOI:** 10.1101/598482

**Authors:** Rosemary Hartman, Stacy Sherman, Dave Contreras, Alison Furler, Ryan Kok

## Abstract

Restored tidal wetlands may provide important food web support for at-risk fish species in the Sacramento-San Joaquin Delta (Delta) of California, including Delta Smelt (*Hypomesus transpacificus*) and Chinook Salmon (*Oncorhynchus tshawytscha*). Since many tidal wetland restoration projects are planned or have recently been constructed in the Delta, understanding the diversity and variability of wetland invertebrates that are fish prey items is of increasing importance. During this study, two different invertebrate sampling techniques were tested (leaf packs and sweep nets) in four habitat types within three different wetland sites to evaluate which sampling technique provided the most reliable metric of invertebrate abundance and community composition. Sweep nets provided a better measure of fish food availability than leaf packs and were better able to differentiate between habitat types. Generalized linear models showed submerged and floating vegetation had higher abundance and species richness than channel habitats or emergent vegetation. Permutational multivariate analysis of variance showed significantly different communities of invertebrates in different habitat types and in different wetlands, and point-biserial correlation coefficients found a greater number of mobile taxa associated with sweep nets. There were more taxa associated with vegetated habitats than channel habitats, and one region had more taxa associated with it than the other two regions. These results suggest that restoration sites that contain multiple habitat types may enhance fish invertebrate prey diversity and resilience. However, the effect of habitat diversity must be monitored as restoration sites develop to assess actual benefits to at-risk fish species.

## Introduction

Tidal wetlands provide an important source of productivity to many estuaries worldwide, subsidizing the surrounding open-water areas with vascular plant detritus, phytoplankton, zooplankton, and nekton biomass [1–5]. Productive freshwater tidal wetlands dominated the landscape of California’s Sacramento-San Joaquin Delta (Delta) prior to the Gold Rush, but, by 1930, the vast majority of wetlands were reclaimed, primarily for agriculture [6] (see Fig 1A). While there is currently no quantitative estimate of the impact of wetland loss on aquatic primary productivity, production most likely declined drastically post-reclamation [7].

**Figure 1.**
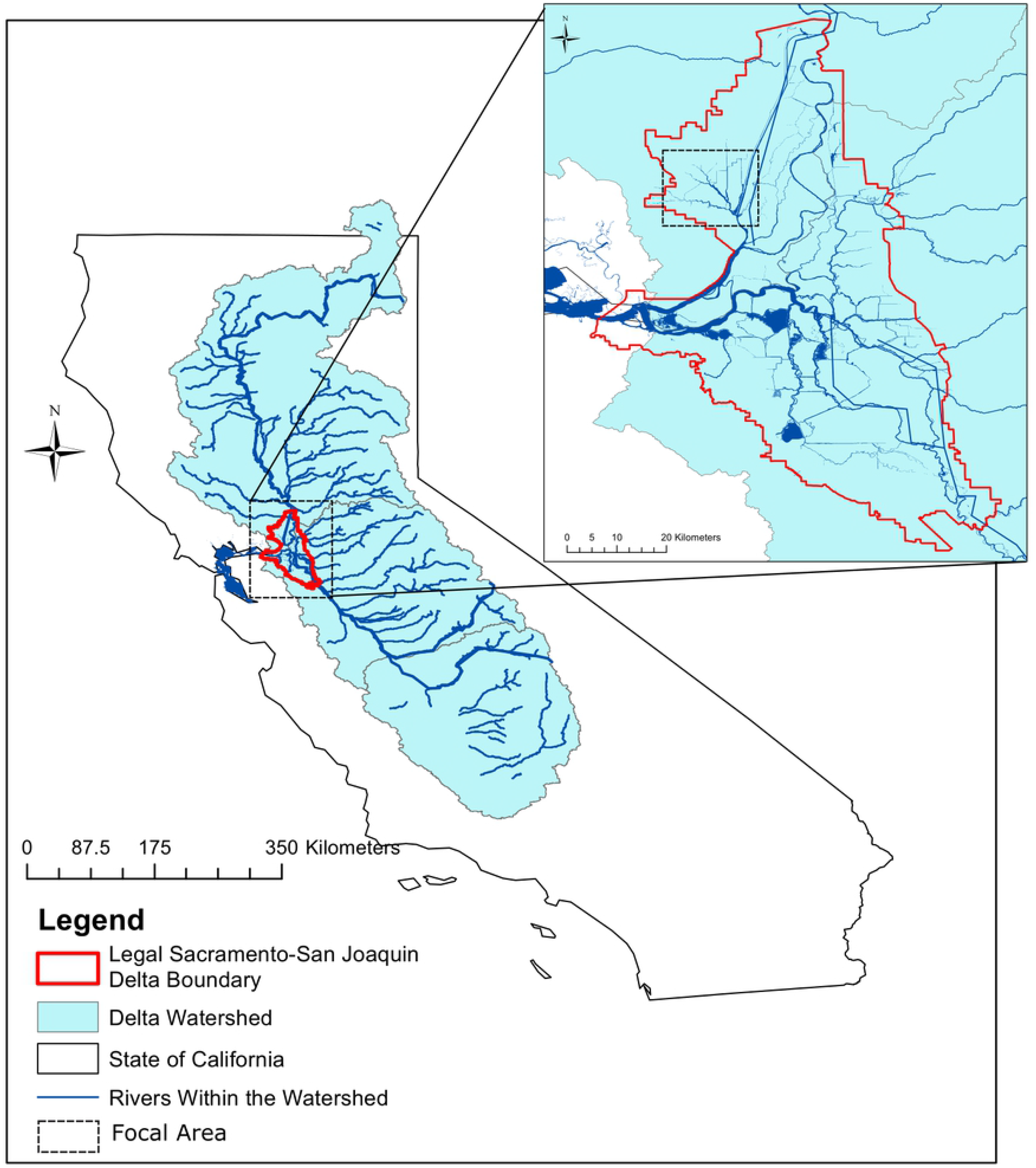
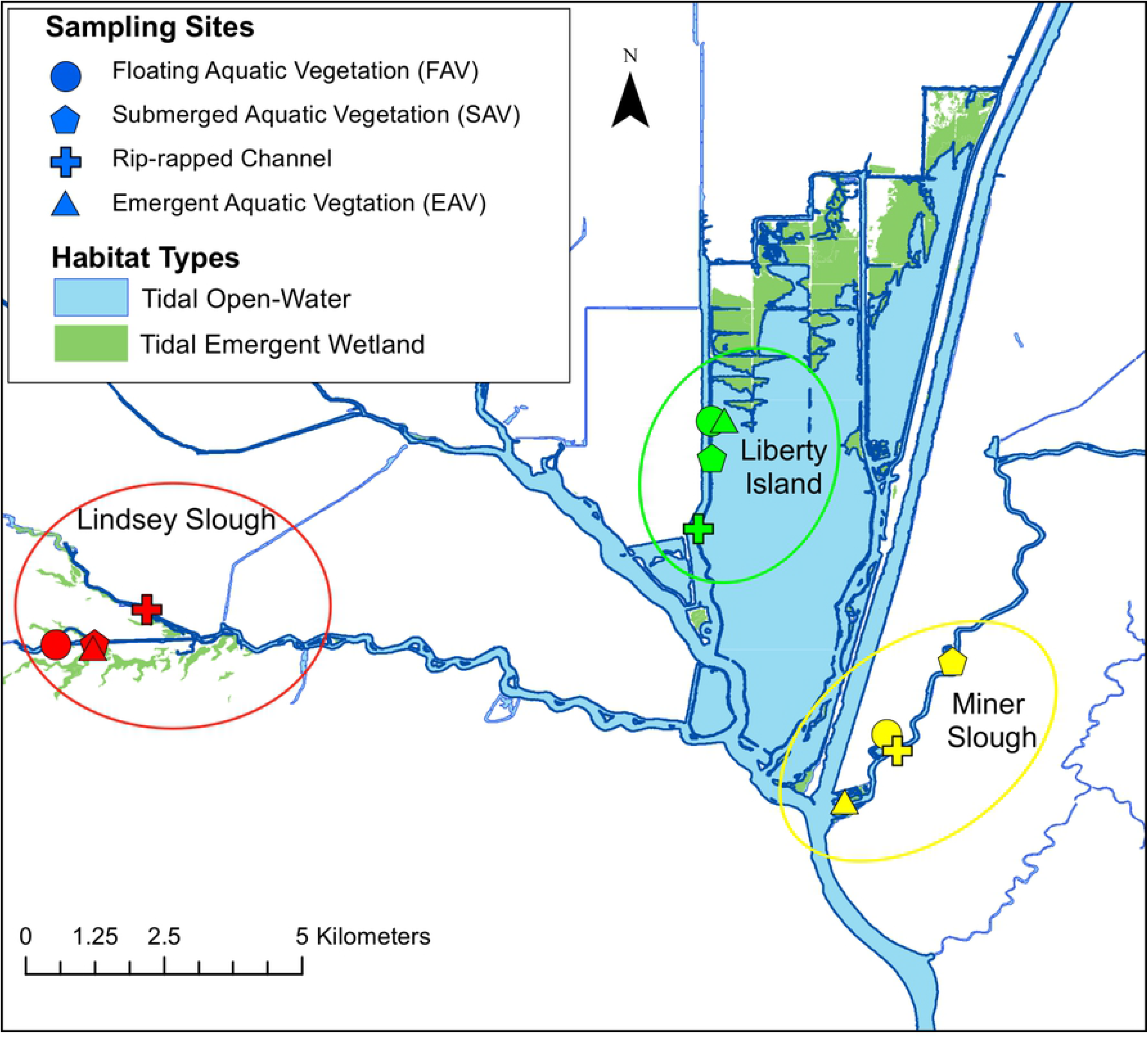
**A)** Map of California (USA) with the San Francisco Bay-Delta watershed. The inset is a finer-scale map of the Legal Delta, with the focal area in the Cache Slough Complex outlined with a dashed box. **B)** The Cache Slough complex, with sampling regions circled. Each region contained four sampling sites with different habitat types: SAV, FAV, EAV, and channel.

Restoration of tidal wetland habitat in the Delta may increase overall primary and secondary production, and thus multiple regulatory mandates now include tidal wetland restoration as part of the recovery effort for several threatened and endangered native fish species, including Chinook Salmon (*Oncorhynchus tshawytscha*) and Delta Smelt (*Hypomesus transpacificus*) [8–11]. Restoration sites of varying size and type are being planned in various locations throughout the Delta and neighboring Suisun Marsh. However, there is a lack of understanding of how to quantify invertebrate abundance within wetlands, and limited knowledge on how invertebrate production varies within and between wetlands.

The current Delta ecosystem is dominated by deep channels, steep, rip-rapped levee banks, and tidal lakes [6]. The aim of wetland restoration is to create a more complex mosaic of habitat types that are believed to be beneficial as fish habitat and for invertebrate production. This mosaic includes shallow, subtidal habitat that provides high phytoplankton production [12] and directly increases pelagic secondary production [13]. Intertidal zones dominated by emergent vegetation also provide a subsidy of detrital carbon and a substrate for epiphytic algae and invertebrates [3, 14, 15]. Wetlands contain complex tidal channel networks that allow fish to access the food produced in the intertidal zone, and provide refuge from predation [16–19]. These diverse habitat types may support unique invertebrate communities, which must be quantified in order to show the effectiveness of wetland restoration.

Some habitat types are not directly targeted by wetland restoration, but often occur in restored habitats and may also provide valuable invertebrate production. Habitats dominated by invasive submerged aquatic vegetation (SAV), such as Brazilian waterweed (*Egeria densa*), and floating aquatic vegetation (FAV), such as water hyacinth (*Eichhornia crassipes*), have been implicated as negative for native fish in the Delta due to high occupancy by invasive predatory fish [20, 21], and adverse effects on water quality [22, 23]. However, invasive vegetation may also provide substrate for large numbers of epiphytic invertebrates [24–26]. Restoration plans often try to limit the establishment of SAV and FAV, but invasive vegetation will be inevitable at some locations.

Despite the need to assess restoration effectiveness, there is no recognized standard for sampling epibenthic and epiphytic invertebrate biomass in wetlands. Though previous studies have been conducted to determine the most effective collection methods for invertebrates in standing water, there is currently no one method that is broadly agreed upon [27]. Some methods, such as drop-frames, may provide high-quality data, but can be time and cost prohibitive [28]. Other methods, such as Hester-Dendy disk sets, have provided good data in some systems [27] but are highly dependent on abiotic variables, and early trials in the Delta yielded very low catch [29]. Invertebrate sampling methods that have been employed in other wetlands often prioritize diversity and presence of sensitive species (as an index of biotic integrity) rather than biomass or abundance [e.g. 30, 31, 32]. Furthermore, wetland conditions, such as tidal influence, topographic complexity, and vegetation structure, vary between bioregions. Because of this, a method that has been proven effective in the sawgrass prairies of the Everglades (for example) may not work in the tule (*Schoenoplectus spp*.) marshes of the Delta [33]. To determine whether wetland restoration provides increased food for at-risk fishes, studies should evaluate differences in invertebrate density and biomass over time and between habitats. Wetlands are a mosaic of different habitat types, so methods to measure invertebrate biomass must work consistently across all habitats.

We hypothesized that a passive colonization substrate sampler, which could be deployed in the same way in multiple habitats, would provide a more controlled measure of invertebrate abundance than active methods (such as nets or benthic grabs). However, we were unsure whether the potential reduction in variability would be worth the increased effort required by the multiple trips to the sampling locations necessary for colonization substrates. Therefore, after an extensive literature review, we conducted preliminary trials of multiple passive methods (Hester-Dendy disk sets, mesh scrubbers, and leaf packs), and compared them against multiple active methods (sweep nets, Marklund samplers, throw traps) to determine which were feasible for a study with higher replication (full results available in [29]). Hester-Dendy disk sets and mesh scrubbers had very low catch, resulting in low power for comparing community composition. Throw traps, which have been very successful in other wetlands [28, 33], did not work well in the tall, thick tule marshes of the Delta. Marklund samplers were difficult to use effectively and had few comparable studies [34]. Of these methods, sweep nets were the most effective active method and leaf packs were the most effective passive method.

In this study, we followed up on our previous trials and evaluated leaf packs, colonization substrates made of standardized bundles of the dominant vegetation left in the wetland for several weeks (as used in [35]), and sweep nets (d-frame nets swept through the water several times by hand, as used in [33]) to see which was most effective in describing the diversity and density of invertebrates. Leaf packs are commonly used for stream systems but are also used in wetland and estuarine systems where there is extensive emergent vegetation [35–37]. Sweep nets often capture higher invertebrate species diversity than colonization substrates, though with higher variability in biomass [33].

We had three major research questions in this study:

1. How do leaf packs compare to sweep nets in collecting a sample of the invertebrate community?

a) Which sampling method has higher power to differentiate between habitat types?
b) Which sampling method has higher power to differentiate between wetlands in neighboring sloughs?
2. How do invertebrate communities change across wetland habitat types?
3. How do invertebrate communities change between wetlands in neighboring sloughs?

We hypothesized that leaf-packs would be easier to standardize across habitat types, and thus provide a lower-variance, higher-power method to differentiate between habitat types and wetland sites. We expected relatively large differences in community composition between habitat types, but small differences between wetlands in neighboring sloughs.

## Methods

### Sample Location and Timing

We conducted two intensive bouts of sampling, one on March 16-17, 2016, and one on May 2-3, 2016. Because salmon and smelt are both anadromous and semi-anadromous, respectively, they are not present in the freshwater wetlands year-round. Spring (February-May) is the period of upstream migration of Delta Smelt the peak residence of juvenile salmonids [38–40]. This is not the period of highest amphipod and insect abundance [2, 26], but the salmon and smelt that consume these are present at their highest densities, and are therefore most able to take advantage of the available invertebrates.

All samples were collected in the Cache Slough Complex, an area in the north-east Delta with high freshwater tidal wetland restoration potential because of high native fish density and appropriate intertidal elevations (Fig 1B)[40, 41]. We chose three regions within the Cache Slough Complex to provide a range of wetland habitats: 1) Liberty Island, 2) the downstream end of Miner Slough and its adjacent marshes, and 3) the Lindsey Slough Restoration site. Liberty Island, an island formerly diked for farming, was flooded by an accidental levee breach in 1997 and contains one of the largest emergent tidal marshes in the present-day Delta. Miner Slough is a distributary of the Sacramento River that flows past Prospect Island, a future tidal wetland restoration site [42]. Lastly, Lindsey Slough Restoration site is a dead-end slough with a formerly diked wetland that was restored to tidal action in the fall of 2014. All three sites were in public waterways or land owned by the California Department of Fish and Wildlife (CDFW).

### Description of Habitat Types

In each region, we targeted four habitat types typical of tidal wetlands. Emergent aquatic vegetation (EAV) sampling sites were in dense stands of native tules (*Schoenoplectus* spp). Submerged aquatic vegetation (SAV) sampling took place in dense stands of invasive Brazilian water weed (*Egeria densa*). Floating aquatic vegetation (FAV) sampling was conducted in dense patches of invasive water hyacinth (*Eichhornia crassipes*). Channel sampling occurred in major channels outside the vegetated wetlands, where the banks were reinforced with large concrete chunks and boulders (rip-rap). We planned to collect three samples per habitat type per region per time period, for a total of 18 sweep nets and 18 leaf packs per habitat type. However, SAV and FAV were not present at all sites in March, and not all leaf packs were recovered due to high flows, vandalism, and loss, which resulted in a reduced sample size.

### Description of Sampling Methods

For all invertebrate sampling methods, we used 500 µm mesh nets and sieves to target macroinvertebrates. All samples were preserved in 70% ethanol dyed with rose Bengal. All sampling was conducted under the Interagency Ecological Program Section 7 Biological Opinion issued to the U.S. Bureau of Reclamation by U.S. Fish and Wildlife Service (USFWS) in 1996, and additional amendments directly from the USFWS to CDFW (file number 1-1-96-F-1 and 1-1-98-I-1296). No scientific collection permit was necessary because all staff members were employees of CDFW (Fish and Game Code Section 1001).

#### Sweep nets

We used a 25 cm × 30 cm d-frame net with 500 µm mesh for all sweep net samples. At each site, we collected three replicate samples, at least five meters apart. We adapted the sweep net technique slightly for different habitat types.

**Channel (rip-rap):** Five 1 m sweeps approximately 3 cm above the substrate.
**EAV:** Five 1 m sweeps, scraping the vegetation as much as possible to knock invertebrates off the stems.
**SAV:** Five 1 m sweeps through the thickest growth, collecting vegetation remaining within the net frame at the end of the sweep.
**FAV:** Net was lifted from beneath a clump of *Eichhornia*. Plant material outside of the net frame, and any leaves above the surface of the water, were severed from the sample with shears, leaving roots and associated invertebrates (similar to [43]).

#### Leaf packs

Tules (*Schoenoplectus acutus*) were harvested and dried to constant weight at 60 °C. Each leaf pack consisted of 30 g of dried stems (each approximately 15 cm in length) in a labeled, plastic mesh bag with 1 cm stretch mesh. This mesh was wide enough to allow all macroinvertebrates of interest to enter without allowing the stems to escape. These samplers were suspended in the midst of the vegetation for EAV, FAV, and SAV samples, and pinned on the bottom in channel habitat. Leaf packs were deployed February 2 and collected March 16 and 17, simultaneously with sweep net sampling. A second set of leaf packs were deployed March 16^th^ and 17^th^, and collected at the same time as sweep net sampling May 2 and 3. During collection, leaf packs were surrounded with a 500 μm mesh net to retain macroinvertebrates.

### Laboratory Methods

Preserved invertebrates were counted and identified to varying taxonomic levels, according to their importance in fish diets, then grouped into larger taxonomic groupings (Order or Class) for analysis. All terrestrial invertebrates were grouped into a single “terrestrial” classification. Insects with both aquatic and terrestrial life stages were classified by life stage, with the terrestrial adults grouped into the “terrestrial” classification and the aquatic larvae classified by Order. If less than 400 individuals were present in a sample, the entire sample was identified. If more than 400 individuals were present, or more than four hours were required for processing, the invertebrates were quantitatively sub-sampled using a grid tray.

### Analysis

To determine which sampler type had higher within-site variability, we compared coefficient of variation in total catch between the two groups using Bartlett’s K-squared test. We compared total catch and species richness of the sampler types across habitat types and regions using generalized linear mixed models (GLMMs), with the predictor variables listed in Table 1. We tested the fit of all possible models and their first-order interactions using Akaike’s Information Criterion corrected for small sample sizes (AICc)[44, 45] and selected the top model. Data were log-transformed to meet assumptions of normality and homogeneity of variance. All analyses were performed using Program R version 3.4.1[46], package lme4 [47].

**Table 1.** Predictor variables for explaining observed differences in catch and species richness.

| Variable | Description |
| --- | --- |
| Region | Region of the Cache Slough Complex as shown on Fig 1b (Lindsey Slough, Liberty Island, or Miner Slough) |
| Habitat type | EAV, SAV, FAV, or rip-rapped channel bank |
| Sampler type | Leaf pack or sweep net |
| Month | March or May |
| Error(Site) | Specific sampling location, used as a random effect to prevent pseudoreplication. |

To detect differences in community composition, we calculated the percent relative abundance of each organism in each sample and used non-metric multidimensional scaling (NMDS) to visualize degree of overlap between communities. We used permutational multivariate analysis of variance (PERMANOVA) using the same set of predictor variables as the GLMM to test for statistical differences in community composition (Table 1) with R package vegan [48].

The effect sizes and degrees of freedom calculated from the above analyses were used in a power analysis to determine minimum number of samples for each sampler type necessary to differentiate total catch and community composition between habitat types and regions using R package pwr [49].

To identify which taxa were most strongly associated with certain sampling methods, regions, and habitat types, we calculated the point-biserial correlation coefficient (*r_pb_*) for each taxon, and tested coefficients’ significance using the multipatt function in the R package indicspecies [50]. This statistical technique takes both frequency of occurrence and abundance into account in assigning which taxa are most closely associated with certain variables.

## Results

### Comparison of sweep nets and leaf packs

We recovered 60 of the 72 (83%) deployed leaf packs. Losses during deployment occurred due to vandalism, high flows, or stranding above the high-water mark. In contrast, 66 of the 67 (98.5%) sweep net samples were successfully collected. This gave us a total of 126 samples with 38,032 individual organisms which we divided into 25 taxa (see S1, Data).

Sweep nets had a significantly higher coefficient of variation in total catch than leaf packs (1.53 versus 1.17, Bartlett’s K-squared: 160.9, P value <0.001). The variance in total catch for sweep nets was also higher than leaf packs (Bartlett’s K-squared: 32.39, P value <0.001). Non-homogeneous variances made the data inappropriate for parametric statistics, however, a Kruskal-Wallace test showed that total catch was not significantly different between the two sampler types (test statistic: 0.087, P value: 0.767).

### Total catch and species richness

When ranking all possible models of log-transformed total catch, the highest ranked model (delta AICc > 2) included only region and habitat type (Table 2). There was much less support for models including sample type, month, or interactions between sample type and habitat type. In particular, FAV and SAV samples had higher catch than channel and EAV samples (Fig 2), and catch in Lindsey Slough was higher than in Liberty Island and Miner Slough. Post-hoc power analyses of the top-ranked model of total catch showed that 30 sweep net samples were needed to achieve 80% power at a 0.05 significance level for the overall model, whereas only 13 leaf pack samples were needed.

**Table 2.** Coefficients for top ranked model predicting total invertebrate catch of sweep nets and leaf packs. Only region and habitat type were included in the top model, sampler type and month were not supported. Top Model: log(Catch) ∼ Region + Habitat; Residual standard error: 1.042 on 40 degrees of freedom (DF); Multiple R^2^: 0.4113, Adjusted R^2^: 0.3377; F-statistic: 5.589 on 5 and 40 DF, P value: 0.0005372

| Factor | Estimate | St. Error | t value | P value |
| --- | --- | --- | --- | --- |
| Intercept (Channel, Liberty) | 4.844 | 0.372 | 13.023 | <0.001** |
| Habitat: EAV | 0.271 | 0.425 | 0.638 | 0.527 |
| Habitat: FAV | 1.431 | 0.435 | 3.287 | 0.002** |
| Habitat: SAV | 1.501 | 0.435 | 3.448 | 0.001** |
| Region: Lindsey | 0.934 | 0.375 | 2.493 | 0.017* |
| Region: Miner | 0.044 | 0.382 | 0.115 | 0.91 |

\* significant P value, $P < 0.05$
\*\* highly significant P value, $P < 0.005$

**Figure 2.**
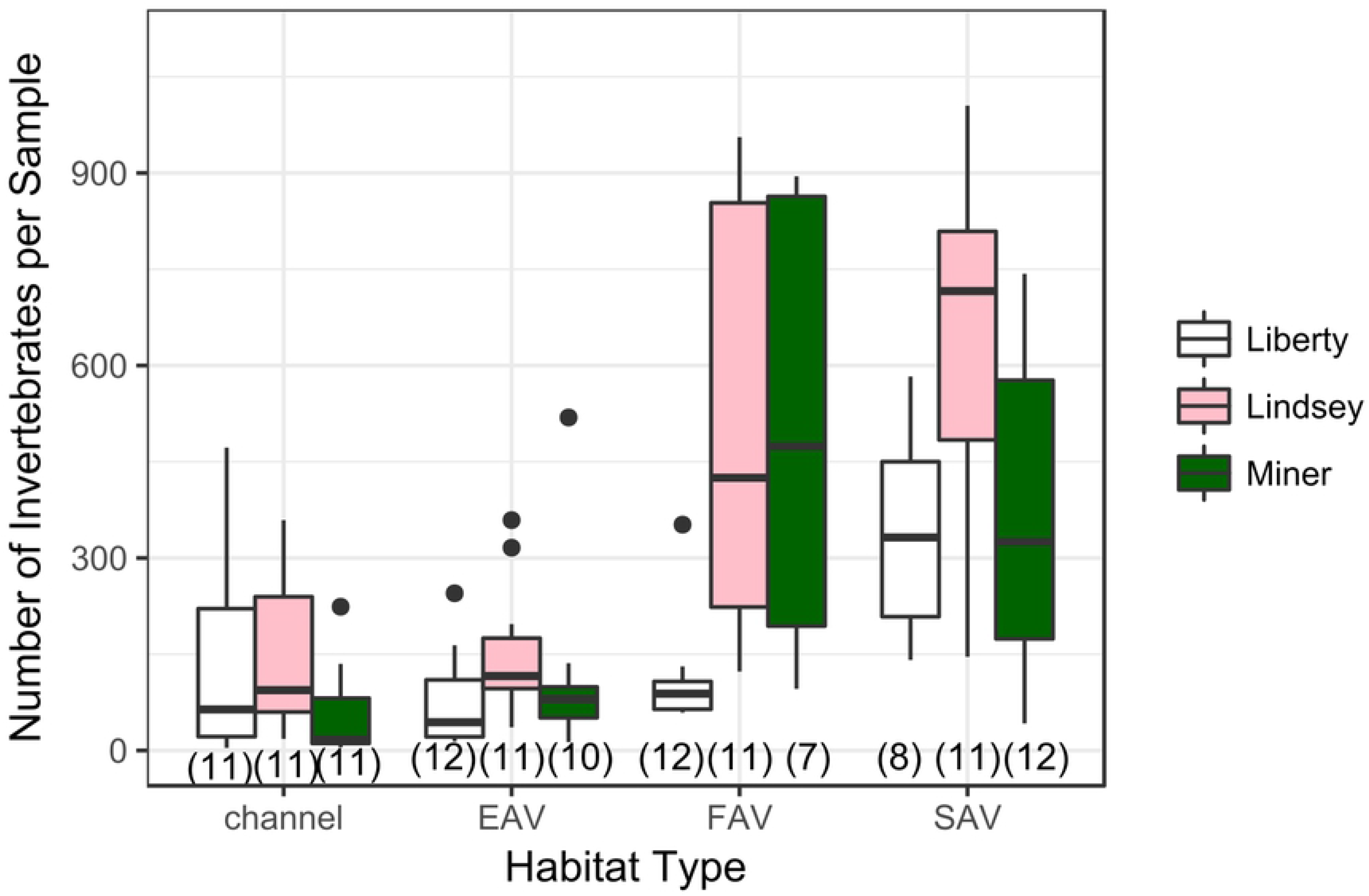
Distribution of total catch in each region and habitat type. Sample size in parentheses below boxes. Four outliers in FAV and SAV samples with catch > 1000 not shown. Models support significantly higher catch in FAV and SAV than in EAV and channel habitats, and significantly higher catch in Lindsey Slough than in Miner Slough or Liberty Island (see Table 2).

When ranking all possible models of species richness, the highest ranked model (delta AICc > 2) included only sample type and habitat type (Table 3). There was much less support for models including region, month, or the interaction between sample type and habitat type. In particular, sweep nets had higher species richness than leaf packs, and SAV and FAV samples had higher richness than channel or EAV samples (Fig 3).

**Table 3.** Coefficients for top ranked model predicting species richness of sweep nets and leaf packs. Only sample type and habitat type were included in the top model; region and month were not supported. Top model: Richness ∼ Habitat + Sampletype, Residual standard error: 5.429 on 41 DF, Multiple R^2^: 0.2915, Adjusted R^2^: 0.2224, F-statistic: 4.217 on 4 and 41 DF, p value: 0.005968

| Factor | Estimate | St. Error | t value | P value |
| --- | --- | --- | --- | --- |
| Intercept (Channel,<br>Leaf pack) | 12.808 | 1.76 | 7.725 | <0.001** |
| Habitat: EAV | 3.083 | 2.216 | 1.391 | 0.172 |
| Habitat: FAV | 6.073 | 2.267 | 2.679 | 0.011* |
| Habitat: SAV | 4.593 | 2.267 | 2.026 | 0.049* |
| Sample type: Sweep net | 4.718 | 1.604 | 2.941 | 0.005* |

\* significant p-value, P < 0.05
\*\* highly significant p-value, P < 0.005

**Figure 3.**
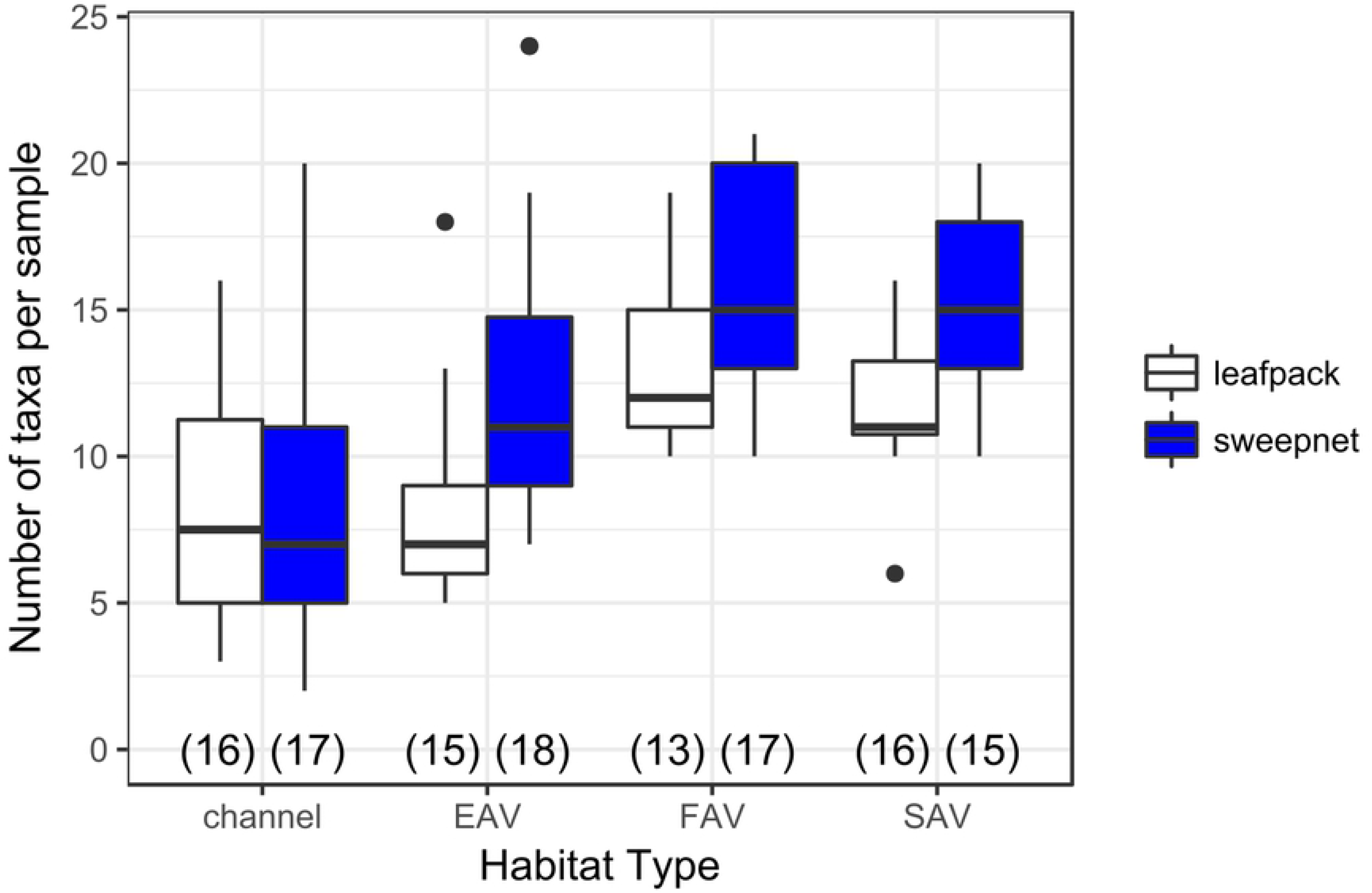
Distribution of species richness for sweep nets and leaf packs in various habitat types. Sample size is in parentheses along the x-axis. Models support significantly higher richness for samples collected with sweep nets, and significantly higher richness for FAV and SAV samples than EAV or channel samples.

### Community composition

Community composition also varied between habitat types and between regions. An overall PERMANOVA showed that there were significant differences between habitat type, sampler type, and region, though not between months. However, pairwise comparisons of PERMANOVA results indicate that sweep nets show significant differences between regions as well as between habitat types, whereas leaf pack samples only had differences between regions and did not show differences between habitat types (Table 4, Fig 4). This can be seen on the NMDS plots, where hulls surrounding habitat types in leaf pack samples have a much higher degree of overlap than hulls surrounding regions (Fig 5A), and the consistent dominance of particular taxonomic groups among habitat types for each region (Fig 4). Sweep net NMDS plots had relatively less overlap between hulls for habitat types (Fig 5B), though habitat explained less of the variation than region (R^2^= 0.16 versus R^2^ = 0.30, Table 4). Post-hoc power analyses of the PERMANOVA results showed that 29 sweep net samples were needed to achieve 80% power at a 0.05 significance level for differences between habitat types, whereas 60 leaf pack samples were needed.

**Table 4.** Results of PERMANOVA performed on the entire data set and on subsets of the dataset using sweep nets only or leaf packs only.

| A)Overall PERMANOVA |  |  |  |  |  |  |
| --- | --- | --- | --- | --- | --- | --- |
| Factor | DF | Sum of Sqs. | Mean Sqs. | F value | R<sup>2</sup> | P value |
| Habitat type | 3 | 1.98 | 0.66 | 3.29 | 0.153 | 0.001** |
| Sample type | 1 | 1.24 | 1.24 | 6.15 | 0.096 | 0.001** |
| Region | 2 | 1.83 | 0.91 | 4.54 | 0.141 | 0.001** |
| Residuals | 39 | 7.84 | 0.20 |  | 0.608 |  |

| B) Leaf packs only |  |  |  |  |  |  |
| --- | --- | --- | --- | --- | --- | --- |
| Factor | DF | Sum of Sqs. | Mean Sqs. | F value | R<sup>2</sup> | P value |
| Habitat Type | 3 | 0.904 | 0.301 | 1.533 | 0.157 | 0.107 |
| Region | 2 | 1.503 | 0.752 | 3.825 | 0.261 | 0.001** |
| Residuals | 17 | 3.341 | 0.196 |  | 0.581 |  |

C) Sweep nets only
| Factor | DF | Sum of Sqs. | Mean Sqs. | F value | R<sup>2</sup> | P value |
| --- | --- | --- | --- | --- | --- | --- |
| Habitat Type | 2 | 0.9532 | 0.477 | 2.554 | 0.161 | 0.011* |
| Region | 3 | 1.801 | 0.600 | 3.217 | 0.303 | 0.003** |
| Residuals | 17 | 3.172 | 0.187 |  | 0.535 |  |

\* significant p-value, $P < 0.05$
\*\* highly significant p-value, $P < 0.005$

**Figure 4.**
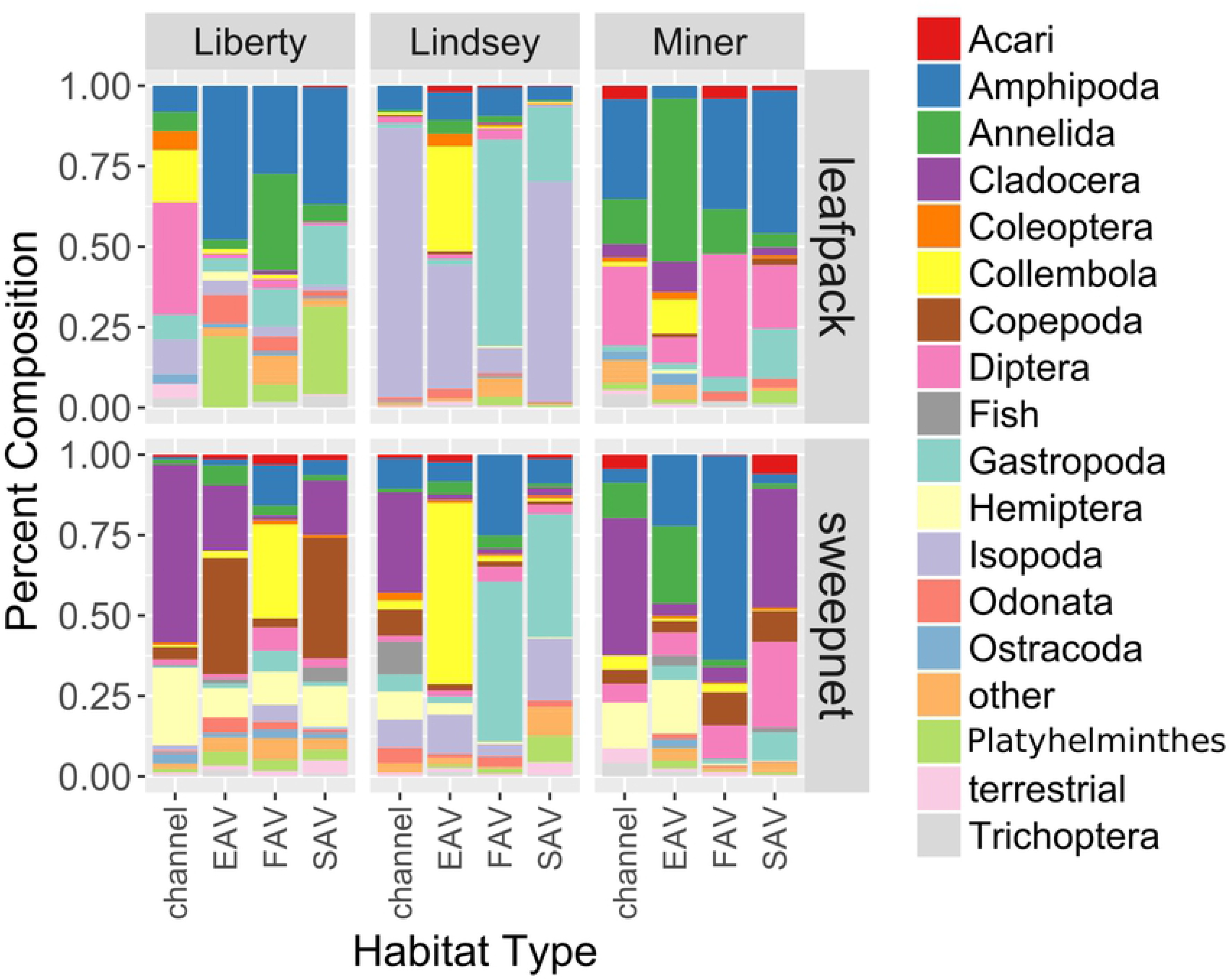
Relative percent composition of major taxa in samples collected with leaf packs and sweep nets in various habitats in the three different regions (Liberty, Lindsey, and Miner). Taxa that made up less than 0.5% of the total catch were combined into the “other” category to simplify the graph. PERMANOVA showed significant differences between habitat types, between regions, and between sample types (Table 4).

**Figure 5.**
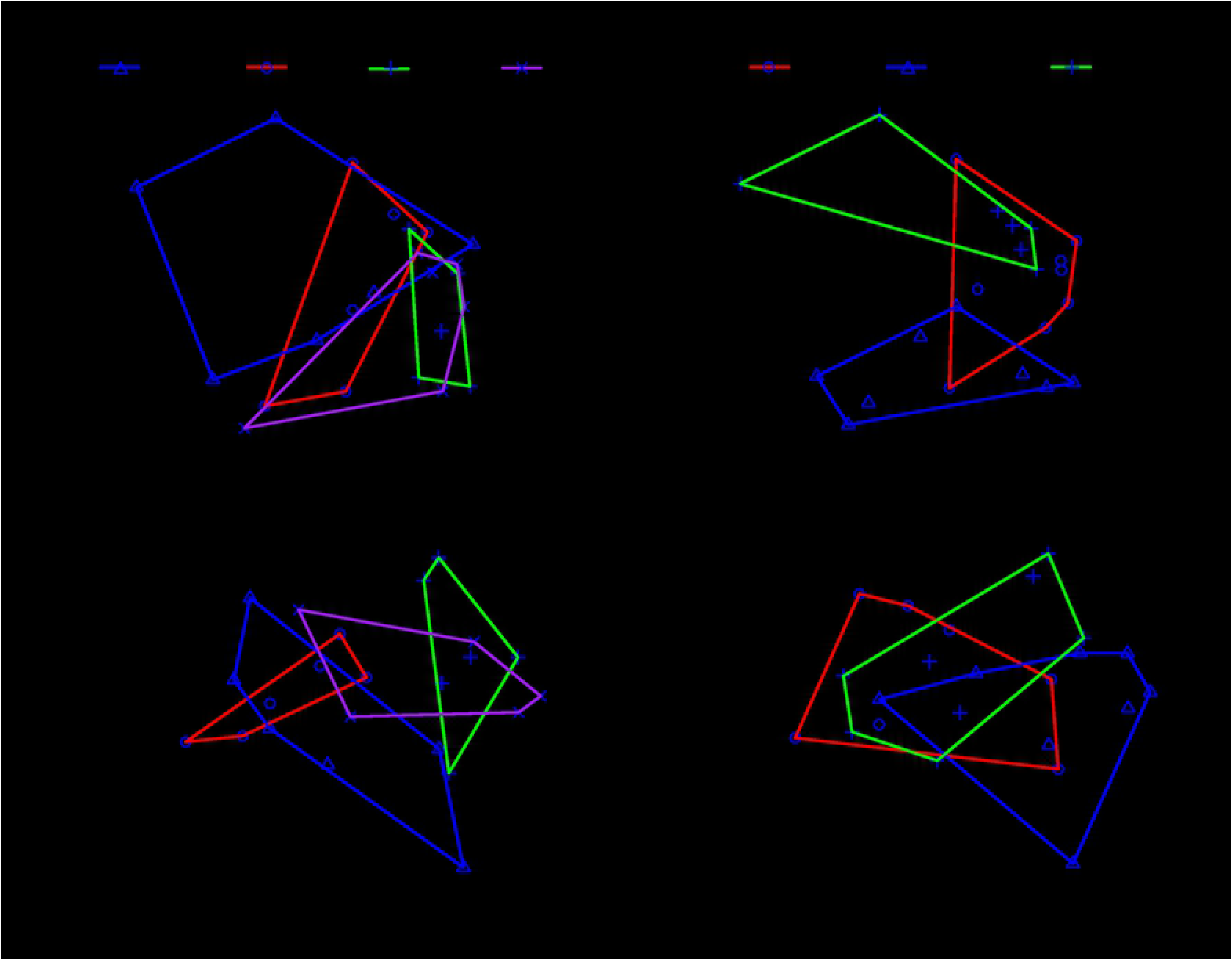
**A)** Non-metric multidimentional scaling plots (NMDS) for all leaf pack data showing grouping by habitat types (replicates summed within sampling sites to avoid pseudoreplication). Stress = 0.159. As can be seen by degree of overlap between hulls, community composition between habitat types was not significantly different (PERMANOVA results: F = 1.56, P = 0.115, R2 = 0.153) **B)** Leaf pack NMDS, with grouping by regions. Community composition between regions was significantly different (PERMANOVA results: F = 3.125, P = 0.004, R2 = .280) **C)** NMDS plots for all sweepnet data showing grouping by habitat types (replicates summed within sampling sites to avoid pseudoreplication). Stress = 0.159. There were significant differences between regions (PERMANOVA F = 3.532, P = 0.002, R2 = 0.211), **D)** Sweet net NMDS showing grouping by regions. There were significant differences between regions (PERMANOVA F = 4.47, P = 0.001, R2 = 0.292)

Analysis of point-biserial correlation coefficients highlight which taxa drove the observed difference in overall community composition between sample types, regions and habitat types (Table 5). Leaf packs were associated with three more sedentary epifaunal taxa, whereas sweep nets were associated with nine highly mobile taxa, including zooplankton and fish (Table 5A). There were four taxa associated with Lindsey Slough, whereas Miner Slough and Liberty Island each had two taxa (Table 5B). There were no taxa associated with channel habitat over the other habitats. One taxon (Collembola) was associated with EAV, three taxa with FAV, and four taxa with SAV (Table 5C).

**Table 5.**
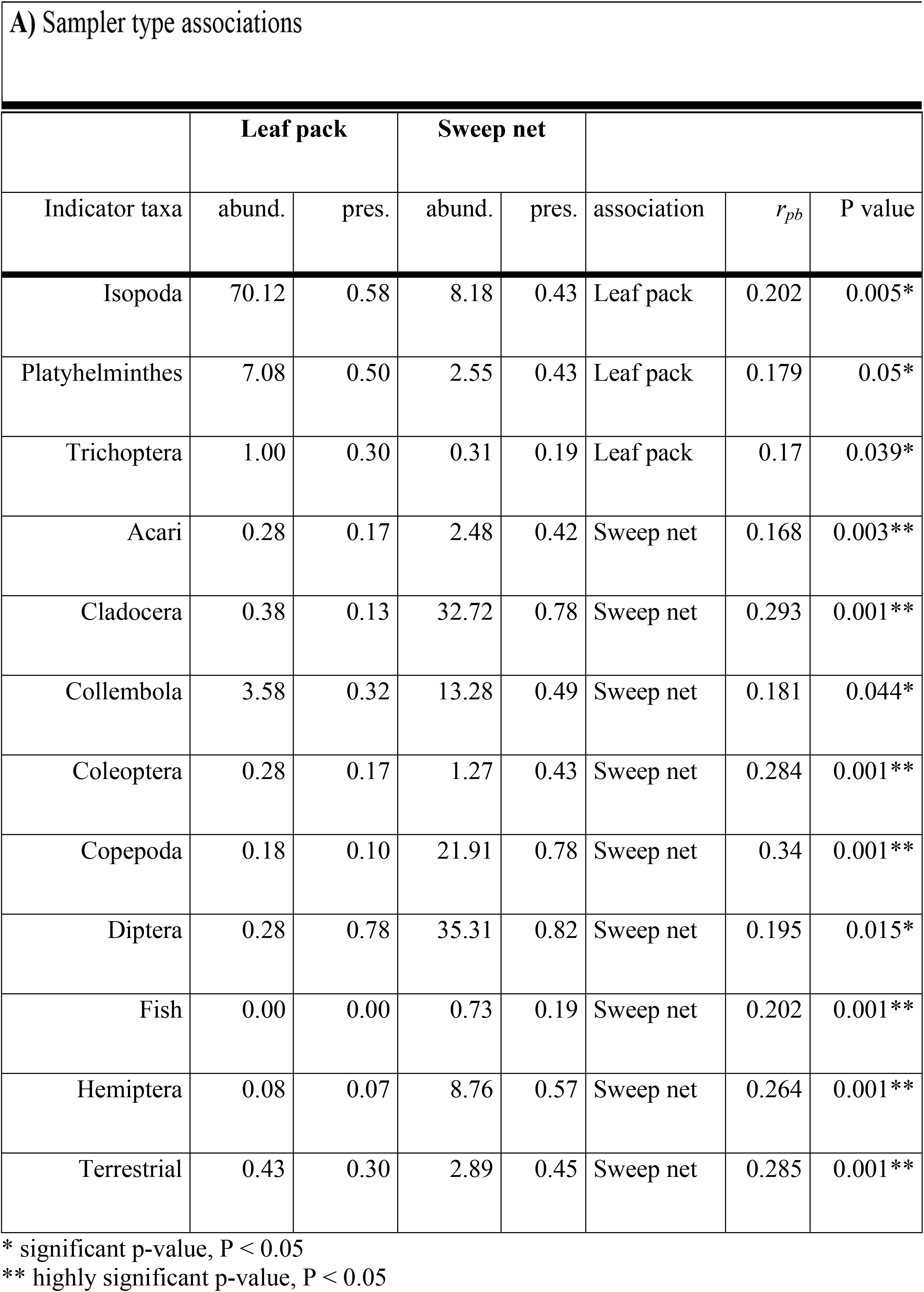
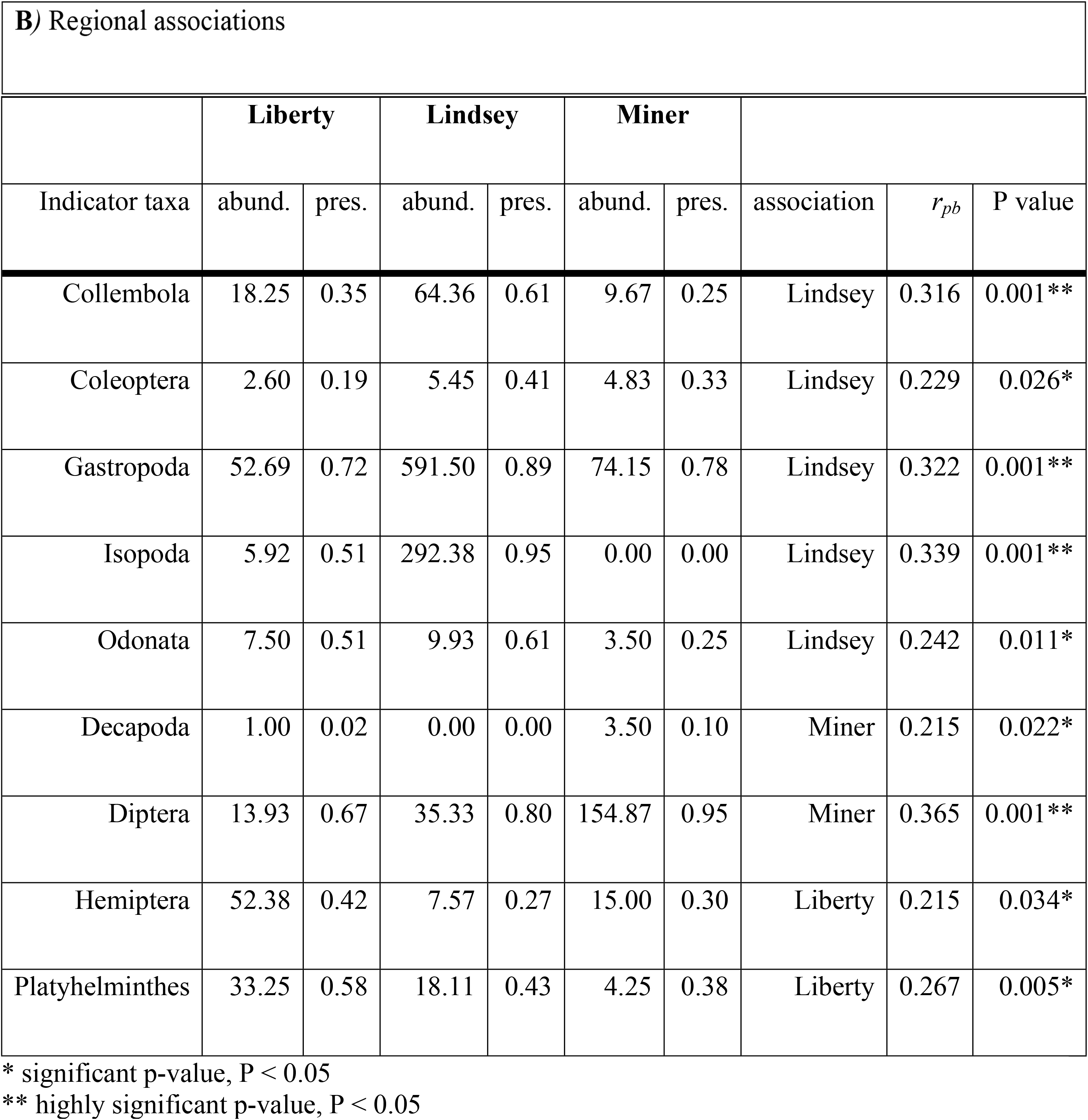
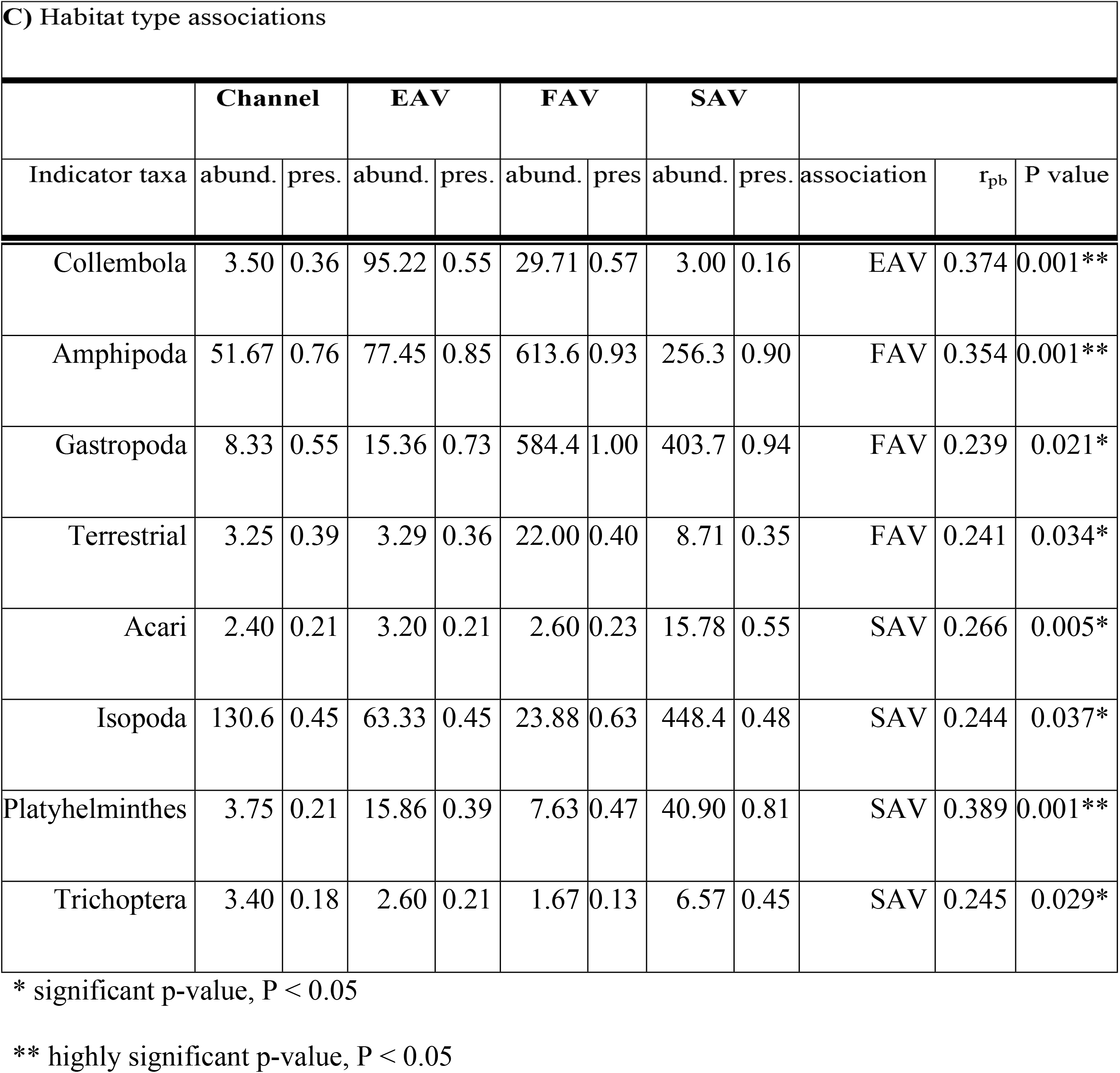
Taxa which were significantly associated with a particular sample type, region, or habitat, mean abundance by group (abund.), proportion of samples where taxa was present (pres.), and statistical significance of the association.

## Discussion

### Choosing sampling methods

We found that sweep nets were more effective and efficient than leaf packs in sampling a variety of shallow water habitats. Sweep nets required a single trip to the field, and did not require assembly ahead of time, making them a considerably lower investment in staff time. While sweep nets had higher variability in total catch than leaf packs, and thus required greater sample sizes to compare catch between groups, they were more cost effective, were less subject to loss or vandalism, were better able to distinguish between habitat types (Fig 5), and had higher species richness (Fig 3, Table 3). They also required fewer samples than leaf packs to characterize community composition. Within the taxa captured by leaf packs, only two taxa were found to be more strongly associated with leaf packs than sweep nets, whereas nine taxa were more strongly associated with sweep nets (Table 5A). Therefore, few taxa will be missed by choosing sweep nets over leaf packs. Furthermore, the insects and Cladocera associated with sweep nets are considered highly important for salmonid diets, whereas the Isopoda and Platyhelminthes associated with leaf packs rarely occur in at-risk fish diets [51, 52].

Our results are consistent with research findings from other areas in which active methods, such as sweep nets, gave a more accurate view of community composition than substrate colonization traps [27, 53]. Sweep nets have also been found to better differentiate between habitat types within a wetland than other sampler types [54].

Because higher sample size is necessary to describe differences in invertebrate density and biomass than to describe differences in diversity (as suggested by [55]), there may be some situations where sweep nets are too highly variable to allow differentiation between regions. In this case, leaf packs may be a valuable alternate sampling method, since they have been used effectively to evaluate wetland restoration in other areas [35]. Furthermore, passive samplers may be more sensitive to different stressors than active methods [27] and may be more appropriate to answer other questions besides production of fish food. However, leaf packs should only be used in emergent vegetation where they most accurately replicate the surrounding habitat and are least likely to be lost.

### Comparing Diversity Across Habitat Types

With both sampling methods combined, we gained a better understanding of how invertebrate communities vary across freshwater wetland habitat types, and how each habitat may be valuable for fish in restored tidal wetland sites.

#### Emergent aquatic vegetation

Emergent vegetation was once the dominant habitat type in the Delta [6], so restoration of this habitat type may provide the best resources for native species. We found a wide variety of taxa in EAV, including Diptera, Hemiptera, and Amphipoda (Fig 4). Collembola were strongly associated with this habitat type, more so than any other habitat (Table 5C). Previous research in the Delta has focused on pelagic invertebrates and open water fish habitat, so there are few local studies of EAV communities with which to compare our results. However, two studies using fall-out traps and neuston tows near EAV in the Delta also found high abundances of Collembola and Diptera [2, 56].

Diptera, Hemiptera, Amphipoda, and Collembola are all important components of fish diets in other estuaries [57–59]. Studies of fish diets from vegetated tidal wetlands in the Delta are scarce, but Sommer et al. [51] found that salmon on the nearby Yolo Bypass floodplain derived the majority of their diets from chironomid midges (Diptera), which are plentiful in EAV (Fig 4). Delta Smelt also appear to consume more insects and amphipods when captured in areas with more EAV [60, 61]. The lack of comparable studies highlights the need for increased research and monitoring in areas of EAV adjacent to future restoration sites.

#### Floating aquatic vegetation

Invasive FAV is actively controlled in the Delta, and many studies have documented the negative impact of *Eichhornia* on water chemistry, water flow, and boat traffic (as reviewed in [23]). However, FAV’s effect on the invertebrate community is understudied. We found a high abundance of invertebrates (Table 2, Fig 2), and found strong associations between FAV and terrestrial invertebrates (Table 5C). We also found strong associations for Amphipoda and Gastropoda, and high abundances of Diptera larvae, similar to other studies of *Eichhornia* in the Delta [25, 43].

While floating vegetation is not considered good habitat for native fishes, the invertebrates we found in this habitat may be an important resource. Terrestrial invertebrates are often an important component of salmonid diets [58], and Amphipoda and Diptera provide particularly high-energy food for at-risk fishes [59]. The benefits of these fish-food invertebrates may help offset the water quality problems associated with *Eichhornia*; however, native species of floating vegetation, such as *Hydrocotyle*, often have a higher overall diversity of invertebrates and higher proportion of native invertebrates [25].

#### Submerged aquatic vegetation

Like FAV, SAV is actively controlled, but few researchers have assessed its invertebrate communities. We found strong associations between SAV and Isopoda, Platyhelminthes, Acari, and Trichoptera (Table 5C), and also found high abundances of Amphipoda, and Diptera (Fig 4). Boyer et al. [24] compared invertebrate communities on *Egeria densa* and *Stuckenia spp*. in Suisun Bay and the western Delta, finding similarly high abundances of Amphipoda, Isopoda, Gastropoda, and Diptera [24]. A similar study by Young et al. [26] in the central Delta that looked at a wider variety of SAV species also found catches dominated by Amphipoda, Diptera, and Gastropoda, though found fewer Isopoda.

Amphipoda and Diptera may be particularly important in salmonid diets [51, 58], so SAV may provide a source of fish food, if the fishes can access it. However, recent expansions of *Egeria* and other invasive SAV in the Delta have been linked to reduced turbidity and increased habitat for non-native piscivores [20–22]. Fish often have decreased foraging success in vegetated habitats [20, 62] and *Egeria densa* decreases foraging success more than other species of SAV [63]. This may decrease non-native piscivores’ ability to prey on native species, but also may decrease the foraging success of native species. Whether increased food production in SAV will offset the negative impacts remains to be seen.

#### Channel habitat

Channel habitat, dominated by rip-rapped banks, is the dominant habitat type in the present Delta ecosystem [6]. Restoration aims to decrease the area of reinforced banks, replacing them with shallow, sloping banks, setback levees, and vegetated benches [64]. There was no significant difference in total catch of invertebrates between channel habitat and EAV (Fig 2, Table 2); however, there was lower taxonomic richness in channel habitat, and the habitats had different community compositions (Fig 3 and 4). We found a relatively high proportion of zooplankton, particularly Copepoda and Cladocera, in these samples (Fig 4), but no taxa uniquely associated with channel habitat (Table 5).

Copepoda and Cladocera are commonly found in salmon and smelt diets [51, 52]. However, these organisms are, on average, smaller and less nutritious than the amphipods common in emergent vegetation [59]. Because all the taxa present in the channel were also found in the other habitat types in similar abundances, this habitat does not appear to provide unique resources for fish. Furthermore, channel habitats are often characterized by rip-rapped banks and man-made structures where predatory fish, such as Striped Bass congregate [65].

### Invertebrate Diversity Across Regions

There were strong differences in abundance and community composition among the three sampling regions (Fig 2, 3, 4), despite all being within ten miles of each other, and in similar sized sloughs. This is in contrast to Simenstad et al. [66], who found relatively small differences in invertebrate communities between sites in the Delta that were much more widely distributed. Thompson et al. [67], found benthic communities in the Delta could be categorized into at least three clusters, though these were based on habitat characteristics (sediment type, vegetation, depth), rather than location *per se*. Other studies of shallow-water habitat in the Delta have found significant differences in phytoplankton and benthic invertebrate biomass that can be traced to tidal transport processes, basin geometry, and benthic substrate [2, 68].

In our study, flow from the Sacramento River greatly influenced water quality on Miner Slough, providing lower turbidity and higher flows than the other two sites. This site provided more decapod crustaceans and Diptera larvae (Table 5b). As a backwater slough, Lindsey Slough had lower flows and longer residence time, characteristics that have been implicated in increased zooplankton productivity, which may also apply to other invertebrates [69]. Lindsey Slough had strong associations with Isopoda, Odonata, Coleoptera, and Collembola (Table 5b). Liberty Island’s primary water source is the Yolo Bypass floodplain, which may be a higher source of productivity than riverine water [70]. This site was particularly important for Platyhelminthes and Hemiptera (Table 5b). Liberty Island also has a much larger area of open water adjacent to our sampling sites than the other two regions, with the potential for increased wind-waves and phytoplankton productivity, which may impact invertebrate abundance [71]. Our observed regional differences may be due to habitat factors not included in our models, such as water velocity, water source, substrate type, and average depth. Further research is necessary to tease apart potential causes for these differences.

### Restoration Implications

We found significant differences in macroinvertebrate communities that may be traced to habitat heterogeneity. This implies that constructing a diverse range of habitats during tidal wetland restoration may increase the variety of invertebrates produced on the site. Fish have dietary preferences, but many fishes shift their diets with the abundance of local resources [72]. For example, Mississippi Silversides collected on Liberty Island were found to consume more amphipods in open water and more insects in vegetated habitat [60]. Delta Smelt collected in deep channels were found to consume less than 5 % amphipods (by weight), and not enough insects to report [52]. However, smelt collected on Liberty Island, where more vegetated habitat is available, consumed 14 % amphipods and 15 % insects [60]. An increase in invertebrates associated with vegetation as part of wetland restoration may help ameliorate declines in the pelagic zooplankton that often make up the majority of smelt diets [73, 74].

A higher diversity of invertebrates may increase resiliency of at-risk fishes and the ecosystem as a whole. There are multiple ways a diverse ecosystem may respond to changes in species composition [75], but there is broad consensus that decreased diversity will decrease food web stability and resilience to change [76, 77]. While restoring a diversity of tidal wetland habitats alone is unlikely to reverse the declines of at-risk fishes, the increase in food web stability provided by wetlands may increase their resilience to other stressors and future disturbances [13, 78]. Differences in diversity and species composition among regions and habitats stress the importance of restoring habitat diversity. Within a restoration site, construction of multiple habitat types may be more beneficial than a single habitat type that is believed to be most important to at-risk species at the time the site is built. We found major differences in communities across sloughs that were relatively close together, and restoration sites spread across the Delta have the potential to provide even higher differences in diversity. However, connectivity between these restoration sites will be essential for migratory species to access all of these diverse resources [13, 64], and for long-term population and community stability [5, 79].

Surprisingly, we found no significant differences in invertebrates between the two time periods (March versus May). This suggests that invertebrate production may remain somewhat constant throughout the spring, however further sample points are required before any conclusions can be drawn. Due to strong seasonal patterns in invertebrate communities found by other studies [2, 36, 80], further research will be necessary to characterize year-round invertebrate communities.

## Conclusion

Invertebrate community composition was highly variable within and between Delta wetlands. Sweep nets provided a simple, efficient way to sample the invertebrate community, and demonstrated that different regions and habitat types supported different groups of organisms. Measuring invertebrate abundance across all habitat types within a wetland will allow managers to evaluate the effectiveness of their restoration projects in providing food for at-risk fishes.

Wetland restoration can benefit from incorporating multiple habitat types in each project to develop a diverse community of invertebrates. Furthermore, restoration projects within different sloughs may have different benefits, and a variety of restoration projects may provide greatest resilience for the aquatic food web and the at-risk fishes it supports.

## Acknowledgements

The authors would like to thank the Fish Restoration Monitoring field and lab crew: J. Mauldin, M. Siepert, C. Hagen, B. Wang, K. Griffiths, S. Lee, and many volunteers. Thanks to V. Tobias and S. Khanna for providing statistics and data visualization support. Thanks to the CDFW Bay Study laboratory staff for help processing invertebrate samples and to S. Lawler, E. Wells, T. Bippus, L. Warkentin, E. Donely-Marineau and W. Fields for help in invertebrate identification. In addition, many thanks to Alice Low and the Interagency Ecological Program Tidal Wetland Monitoring Project Work Team for their guidance and assistance in this project

## Author Contributions

Rosemary Hartman - Conceptualization, Data curation, formal analysis, Investigation, Methodology, Writing – Original draft

Stacy Sherman - Conceptualization, formal analysis, Investigation, Methodology, Project Administration, Writing – review and editing

Dave Contreras – Conceptualization, formal analysis, Investigation, Methodology, Writing – review and editing

Alison Furler –Investigation, Methodology, Writing – review and editing Ryan Kok –Investigation, Methodology, Writing – review and editing

## Funding

This work was funded by the California Department of Water Resources State Water Project and Fish Restoration Program, CDFW contract number R1630002.

## Competing interests

None of the authors have any competing interests to report.

## Supporting Information Caption

S1. Data. Catch of each taxonomic group of invertebrates used for the anlaysis of this data.

